# ASTF1, an AP2/ERF-family transcription factor and ortholog of cultivated tomato LEAFLESS, is required for acylsugar biosynthesis

**DOI:** 10.1101/2022.04.04.487036

**Authors:** Sabyasachi Mandal, Yohannes H. Rezenom, Thomas D. McKnight

**Affiliations:** Department of Biology, Texas A&M University, College Station, Texas 77843, USA; Department of Chemistry, Texas A&M University, College Station, Texas 77843, USA

## Abstract

Acylsugars, specialized metabolites produced by solanaceous trichomes, provide protection against biotic and abiotic stresses. Here, we report ACYLSUGAR TRANSCRIPTION FACTOR1 (ASTF1/Sopen05g008450; AP2/ERF-family member) positively regulates acylsugar biosynthesis. Virus-induced gene silencing (VIGS) of *ASTF1* in *Solanum pennellii* reduced acylsugar production by 65%. Most acylsugar (and several flavonoid) metabolic genes were downregulated in *ASTF1*-silenced plants, and these genes showed strong co-expression with *ASTF1*. In promoters of potential ASTF1-targets, we identified three enriched motifs, and one motif showed similarity with binding sites of other AP2/ERFs. Phylogenetic analysis and data mining indicated trichome-enriched expression of *ASTF1* orthologs in several acylsugar-producing solanaceous species, suggesting a conserved role in acylsugar biosynthesis. This was supported by VIGS of *ASTF1* orthologs in *Nicotiana benthamiana*. Broader phylogenetic analysis revealed relationships among specialized metabolic AP2/ERFs in several asterid species and provided clues about evolutionary emergence of acylsugar phenotype. Cultivated tomato ortholog (LEAFLESS/Solyc05g013540) has been reported to coordinate leaf initiation with transient expression at incipient primordia, and data mining revealed downregulation of trichome-preferentially-expressed genes, including acylsugar (and flavonoid) metabolic genes, in *leafless* mutants’ shoot apices, indicating remarkable spatiotemporal functional diversity. Our work will pave a way to disentangle acylsugar regulatory network and holds promise for future metabolic engineering of acylsugar production.

## INTRODUCTION

Plants produce a myriad of phylogenetically-restricted specialized metabolites which play important roles in plant-environment interactions. These compounds are widely used as medicines and other valuable phytochemicals. Acylsugars are specialized metabolites secreted through trichomes of many plants in the Solanaceae, which includes important agricultural crops such as tomato (*Solanum lycopersicum*), potato (*S. tuberosum*), eggplant (*S. melongena*), and tobacco (*Nicotiana tabacum*). These compounds have major roles in plant defense against herbivores, insect pests and microbial pathogens (Juvik et al., 1994; Chortyk et al., 1997; Hare, 2005; Weinhold and Baldwin, 2011; Luu et al., 2017). Many acylsugars exhibit stronger insecticidal activity than standard, insecticidal soap (Puterka et al., 2003). Additionally, they protect plants from desiccation (Feng et al., 2021). Metabolic engineering of acylsugar production offers a great potential for increasing acylsugar-mediated plant defense and desiccation tolerance. However, success depends on identifying critical regulatory factors that control structural genes in acylsugar metabolic pathways.

Acylsugars are glucose and sucrose esters of both branched- and straight-chain fatty acids (Kroumova et al., 2016; Moghe et al., 2017), which are derived from branched-chain amino acids (Val, Leu, and Ile) and from acetyl-CoA (fatty acid synthase mediated *de novo* biosynthesis), respectively (Walters and Steffens, 1990; Mandal et al., 2020). ACYLSUGAR ACYLTRANSFERASEs (ASATs) attach these acyl groups (C2-C12 in length) to the sugar moiety, and different species of the Solanaceae have evolved specific sets of ASATs to synthesize acylsugars (Schilmiller et al., 2015; Fan et al., 2016; Moghe et al., 2017; Nadakuduti et al., 2017; Feng et al., 2021). Evolution of catalytic functions of ASATs to combine products of primary metabolism (sucrose and acyl molecules) into specialized metabolic acylsugars was a critical step in the evolutionary emergence of acylsugar biosynthesis. Functionality of the newly evolved ASATs and other acylsugar metabolic proteins required recruitment of transcription factors (TFs) to ensure proper gene expression in trichomes, the presumable site of acylsugar biosynthesis in the Solanaceae. Identification of these TFs has been an open and interesting question in the study of acylsugar metabolism.

Metabolic pathways are tightly controlled, and TFs play important roles in regulating spatial, temporal, and environmental expression of structural genes by binding to *cis*-regulatory elements of target genes. The APETALA2/ETHYLENE RESPONSE FACTOR (AP2/ERF) family is one of the largest families of plant TFs. These proteins are characterized by one or two copies of the conserved AP2 DNA-binding domain of about 60 amino acids. This domain is found predominantly in plants, and it may have been acquired by lateral gene transfer from bacterial and viral endonucleases (Magnani et al., 2004). AP2/ERF TFs have important regulatory roles in a wide variety of biological and physiological processes, such as growth and development, plant-environment interactions, and biosynthesis of metabolites, including specialized metabolites. For example, AP2/ERF members regulate biosynthesis of steroidal glycoalkaloids (SGAs) in *S. lycopersicum* and *S. tuberosum* (Cardenas et al., 2016), nicotine in *N. tabacum* and *N. benthamiana* (Shoji et al., 2010; Todd et al., 2010), terpenoid indole alkaloids (TIAs) in Madagascar periwinkle (*Catharanthus roseus*) (Menke et al., 1999; van der Fits and Memelink, 2000), and artemisinin in trichomes of sweet wormwood (*Artemisia annua*) (Yu et al., 2012; Tan et al., 2015).

Here, we report that one AP2/ERF TF, which we have named ACYLSUGAR TRANSCRIPTION FACTOR1 (ASTF1; Sopen05g008450), is involved in acylsugar biosynthesis in wild tomato *S. pennellii*. Virus-induced gene silencing (VIGS) of *ASTF1* led to a decrease in acylsugar production, which was accompanied by downregulation of multiple acylsugar biosynthetic genes. VIGS of *ASTF1* orthologs in *N. benthamiana* also led to a reduction in acylsugar accumulation. Phylogenetic analysis suggested that orthologs of ASTF1 have a conserved role in acylsugar biosynthesis in solanaceous trichomes, and it appears to have diverged from other specialized metabolic AP2/ERFs. Interestingly, the ortholog of *ASTF1* in *S. lycopersicum* (*LEAFLESS*), which is transiently expressed at incipient leaf primordia, is required for leaf initiation at the shoot apex (Capua and Eshed, 2017), and data mining indicated remarkable spatiotemporal transition in expression and function of this TF.

## RESULTS

We previously identified a network of candidate genes, including six TF genes, involved in acylsugar metabolism in *S. pennellii* (Mandal et al., 2020). These six genes are *Sopen01g037680, Sopen03g036630, Sopen05g008450, Sopen10g031080, Sopen10g032000*, and *Sopen12g021250* (annotated as TFs belonging to TCP family, AP2/ERF family, AP2/ERF family, homeodomain-leucine zipper family, MYB family, and AP2/ERF family, respectively). We focused our initial efforts to characterize TF genes that regulate acylsugar biosynthesis on *Sopen05g008450* (*ASTF1*) for reasons described below.

### Among candidate TF genes, *ASTF1* has highest level of trichome-enriched expression

Many acylsugar biosynthetic genes are expressed in trichome tip cells (Ning et al., 2015; Schilmiller et al., 2015; Fan et al., 2016; Fan et al., 2020). Therefore, we first compared expression levels in isolated stem trichomes with expression levels in underlying tissues of shaved stems for each of the six candidate TF genes. *ASTF1* showed the highest level of trichome-enriched expression (208-fold), based on reverse transcription-quantitative PCR analysis. This expression level is comparable with that of *ASAT* genes (Figure 1). Using reciprocal BLAST, we identified putative orthologs of candidate TF genes in *S. lycopersicum* (Mandal et al., 2020), and according to data of Ning et al. (2015), *Solyc05g013540* (ortholog of *ASTF1*) has the highest level of trichome-enriched expression (252-fold) among candidates.

**Figure 1.**
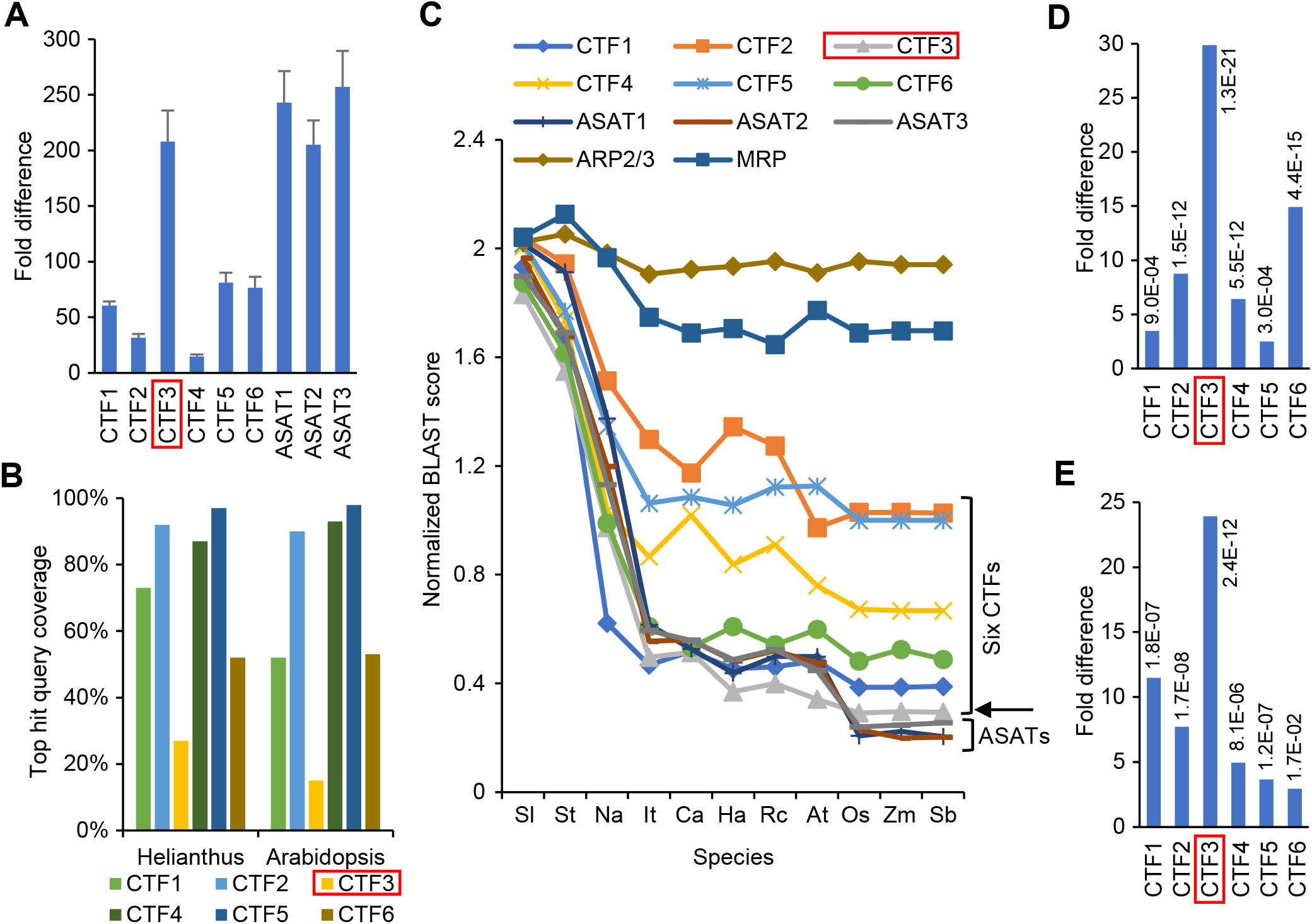
ASTF1 as a candidate acylsugar metabolic TF. Six candidate TFs (CTFs) are: CTF1= Sopen01g037680; CTF2= Sopen03g036630; CTF3= Sopen05g008450 (ASTF1; marked in red boxes); CTF4= Sopen10g031080; CTF5= Sopen10g032000; and CTF6= Sopen12g021250. (A) Expression of six CTF genes in isolated stem trichomes relative to underlying tissues of shaved stems (normalized to one-fold). Error bars indicate standard error (*n* = 5 biological replicates). Orthologs of *S. lycopersicum* trichome-tip-cell-expressed *ASAT* genes (Schilmiller et al., 2015 and Fan et al., 2016) are included for comparisons. (B) BLASTp against *Helianthus annuus* and *Arabidopsis thaliana*. Query coverages for top hits are shown. (C) Extended BLASTp. Total BLAST scores for top hits were normalized by sequence lengths for each sequence. The following species were selected: six asterid species [*S. lycopersicum, S. tuberosum*, and *Nicotiana attenuata* (all belonging to the Solanaceae; Solanales; Lamiid), *Ipomoea triloba* (Convolvulaceae; Solanales; Lamiid), *Coffea arabica* (Rubiaceae; Gentianales; Lamiid), and *Helianthus annuus* (Asteraceae; Asterales; Campanulid)], two rosid species [*Rosa chinensis* (Rosaceae; Rosales; Fabid) and *Arabidopsis thaliana* (Brassicaceae; Brassicales; Malvid)], and three monocot species [*Oryza sativa, Zea mays*, and *Sorghum bicolor* (all belonging to the Poaceae; Poales)]. For comparisons, three ASATs and two conserved sequences [actin-related protein 2/3 complex (ARP2/3; Sopen12g033210) and a mitochondrial ribosomal protein (MRP; Sopen04g006520)] were also included. ASTF1 is indicated with an arrow. (D and E) Differential expression of CTF genes in low- and high-acylsugar-producing accessions of *S. pennellii* (D) and in response to imazapyr treatment (acylsugar inhibitor) (E). False discovery rates are shown next to fold-change values. Data for (D and E) were obtained from Mandal et al. (2020).

### Among candidate TFs, ASTF1 has lowest level of sequence similarity with proteins from distantly related species

Using BLASTp, we identified homologous sequences for each of the six candidate TFs in *S. lycopersicum*, and the top hits showed 100% query coverage for each TF. However, BLASTp against *Helianthus annuus*, a non-solanaceous asterid, revealed that the top hit against ASFT1 covered only 27% of its protein sequence. For the remaining five candidate TFs, coverage ranged from 97% to 52%. In a more distant comparison against *Arabidopsis thaliana*, a rosid, ASTF1 again showed the least amount of coverage with the top hit (only 15%). Coverage for the remaining five candidates ranged from 98% to 52% (Figure 1B). Extended BLASTp analysis with other distantly related species also supported different levels of sequence similarity among candidate TFs, with ASTF1 showing the highest level of sequence divergence (Figure 1C).

### Functional validation of *ASTF1* using virus-induced gene silencing (VIGS)

Its high level of trichome-enriched expression and low level of sequence similarity with proteins of distantly related species strongly suggested a specialized function for ASTF1 in *S. pennellii* trichomes, the site of acylsugar production and secretion. This encouraged us to prioritize *ASTF1* for functional validation. This decision was strongly supported by two other observations. First, *ASTF1* (among the six candidate TF genes) exhibits highest level of differential expression between low- and high-acylsugar-producing accessions of *S. pennellii* (Figure 1D). Second, it also shows strongest repression of gene expression in response to imazapyr treatment, which inhibits acylsugar biosynthesis (Figure 1E).

We performed VIGS using tobacco rattle virus (TRV)-based silencing vectors (Dong et al., 2007), and liquid chromatography–mass spectrometry analysis showed that the total acylsugar amount was reduced by 65% (*P* < 0.0001) in *ASTF1*-VIGS plants (Figure 2, A and B). To corroborate this finding, we performed transesterification reactions to release acyl chains from their sugar moieties and quantified major acyl chains using gas chromatography–mass spectrometry. Compared to a group of control plants (empty TRV vectors), individual acyl chains in VIGS plants were also reduced by 55%-75% (*P* < 0.0001; Figure 2C), which confirmed the involvement of *ASTF1* in acylsugar production. The reduction in acylsugar amount in VIGS plants was not accompanied by a statistically significant decrease in trichome density, and no morphological differences were observed between control and silenced plants (Supplemental Figure 1).

**Figure 2.**
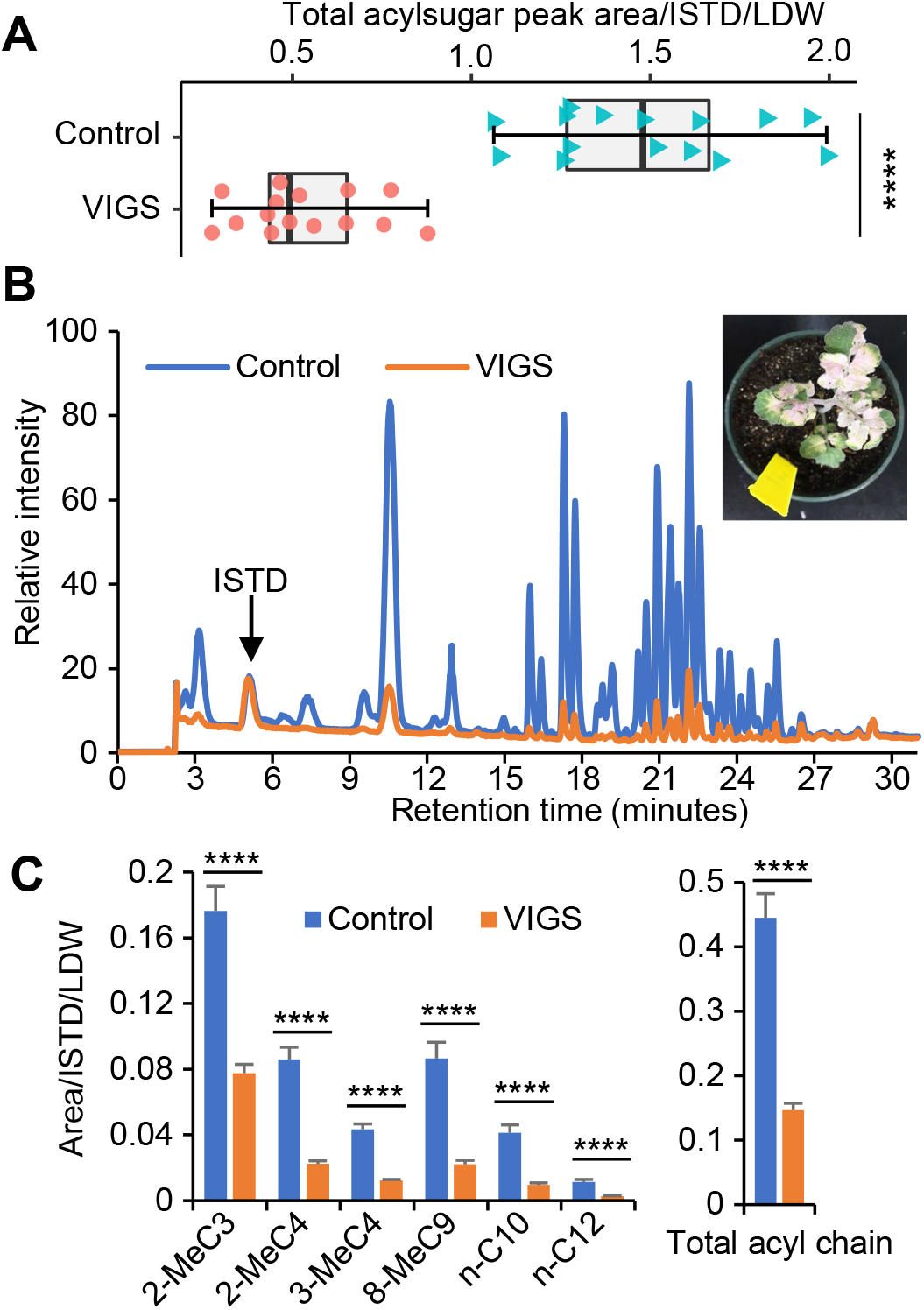
Functional validation of ASTF1. (A) Virus-induced gene silencing (VIGS) of *ASTF1* reduces acylsugar production in *S. pennellii*. Normalized chromatogram peak areas were used to quantify acylsugar amounts (*n* = 15 biological replicates; **** *P* < 0.0001; Welch *t*-test). Normalization was done by dividing total acylsugar peak areas with internal standard (ISTD) area and leaf dry weight (LDW). (B) Representative chromatograms (normalized by internal standard and leaf dry weight) showing acylsugar peaks in control and VIGS plants. The inset shows VIGS of a *phytoene desaturase* gene that was used as a positive control (photobleached phenotype). (C) Acyl chain compositions of acylsugars extracted from control and VIGS plants (major acyl chains are shown). Me= methyl; C3-C12 indicate acyl chain length (for example, 2-MeC3 and *n*-C10 indicate 2-methlpropanoate and *n*-decanoate, respectively). Combined amounts of major acyl chains in control and VIGS plants are shown next to individual acyl chain bar graphs. Error bars indicate standard error (*n* = 15 biological replicates; **** *P* < 0.0001; Welch *t*-test).

### Many known and candidate acylsugar (and trichome flavonoid) metabolic genes are downregulated in VIGS plants

Using RNA-Seq, we identified 88 differentially expressed genes between control and *ASTF1*-VIGS plants (73 downregulated and 15 upregulated in VIGS plants; Supplemental Data Set 1). Gene ontology (GO) terms such as “acyltransferase activity” (GO:0016746), “fatty acid synthase activity” (GO:0004312), and “flavonoid metabolic process” (GO:0009812) were enriched in the list of downregulated genes, whereas GO terms associated with biosynthesis of terpenes (GO:0050551, GO:0050552, and GO:0010333) were enriched among upregulated genes (Supplemental Figure 2). Downregulated genes include most, but not all, genes with known and putative functions in acylsugar metabolism, such as those encoding branched-chain fatty acid metabolic proteins, acyl-activating enzymes, fatty acid synthase components, three ASATs, two of the three candidate ABC transporters, a RUBISCO small subunit, and two of the six candidate TFs (Figure 3).

**Figure 3.**
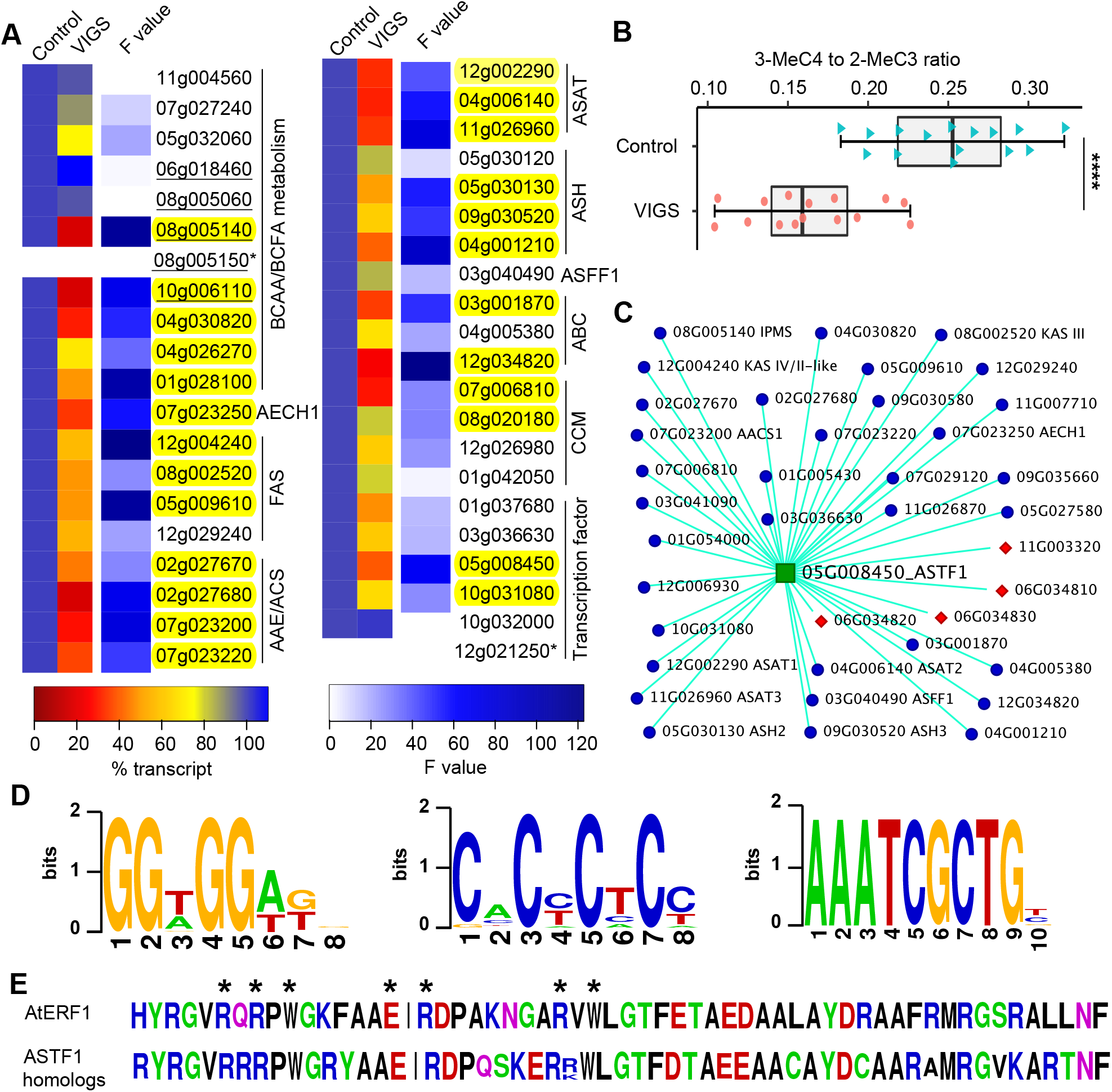
Analysis of ASTF1-target genes. (A) Relative expression levels of known and candidate acylsugar metabolic genes in control (set to 100% transcript) and VIGS plants, based on RNA-Seq analysis (*n* = 5 biological replicates). “*Sopen*” prefix was removed from gene IDs. BCAA= branched-chain amino acid; BCFA= branched-chain fatty acid; FAS= fatty acid synthase components; AAE= ACYL-ACTIVATING ENZYME, ACS= ACYL-CoA SYNTHETASE; AECH1= ACYLSUGAR ENOYL-CoA HYDRATASE 1; ASAT= ACYLSUGAR ACYLTRANSFERASE; ASH= ACYLSUGAR HYDROLASE and the related carboxylesterase Sopen04g001210; ASFF1= ACYLSUCROSE FRUCTOFURANOSIDASE 1; ABC= ABC transporter; CCM= central carbon metabolism. Genes encoding ISOPROPYLMALATE SYNTHASE (IPMS) are underlined. Highlighted genes show statistically significant difference in expression between control and VIGS plants (false discovery rate < 0.05). Two genes (*Sopen08g005150* and *Sopen12g021250*; marked with an asterisk) had low expression levels and were removed before differential gene expression analysis. Individual gene annotation and related information are given in Supplemental Data Set 1. (B) The ratio of 3-methylbutanoate (3-MeC4) to 2-methylpropanoate (2-MeC3). Data points in control and VIGS plants are shown (*n* = 15 biological replicates; **** *P* < 0.0001; Welch *t*-test). (C) Weighted gene correlation network analysis (WGCNA). “Sopen” prefix was removed from gene IDs. Nodes and edges represent genes and intramodular connectivities, respectively. Genes that are strongly co-expressed (based on intramodular connectivities > 0.15) with *ASTF1* are shown. Annotations for known acylsugar metabolic genes are indicated. Red diamonds indicate flavonoid metabolic genes. Detailed WGCNA results are given in Supplemental Data Set 3. (D) Enriched motifs found in promoters of potential ASTF1-target genes. (E) Consensus sequence of the AP2 domain from ASTF1 and 29 homologs in different species of the Solanaceae (listed in Figure 4A). DNA base-contacting amino acids of *Arabidopsis thaliana* ERF1 (AtERF1) (Allen et al., 1998) are indicated with asterisks.

*Sopen11g003320*, a close homolog of a flavonoid (flavonol and flavone) glucosyltransferase (Masada et al., 2009), and three sequential genes on chromosome 6 (*Sopen06g034810, Sopen06g034820*, and *Sopen06g034830*; MYRICETIN *O*-METHYLTRANSFERASE), which are involved in trichome flavonoid biosynthesis (Kim et al., 2014b), were also downregulated in VIGS plants (Supplemental Data Set 1). These *S. pennellii* genes and their putative *S. lycopersicum* orthologs exhibit high levels of trichome-enriched expression (more than 300-fold, according to data of (Ning et al., 2015; Fan et al., 2020)). These results indicate that expression of many genes involved in acylsugar and flavonoid metabolism in trichomes are regulated by ASTF1, either directly or indirectly.

### Some non-acylsugar defense genes are upregulated in VIGS plants

Of the 15 upregulated genes, eight show higher expression levels in a low-acylsugar-producing accession compared to a high-acylsugar-producing accession of *S. pennellii* (Supplemental Data Set 2). These genes include those presumably involved in defense and biosynthesis of other classes of specialized metabolites, such as terpenes and phenolic compounds. For example, *Sopen08g002800* is predicted to encode a patatin-like phospholipase; in *Capsicum annuum*, one such protein is involved in defense signaling (Kim et al., 2014a). Similarly, seven other upregulated genes’ putative orthologs in *S. lycopersicum* have been reported to be involved in plant defense responses. These are *Solyc08g008310* (Iberkleid et al., 2015), *Solyc12g010020* (Pautot et al., 1993; Schimmel et al., 2018), *Solyc02g067750* (Schimmel et al., 2018; Zhang et al., 2020), *Solyc09g084490* (Schimmel et al., 2018; Padmanabhan et al., 2019), *Solyc08g076980* (Schimmel et al., 2018), *Solyc04g077190* (Balestrini et al., 2019), and *Solyc01g067460* (Padmanabhan et al., 2021). In fact, four of these *S. lycopersicum* genes were reported in a single study involving plant-feeding mites (Schimmel et al., 2018). These results corroborate our previous finding that expressions of some non-acylsugar defense genes are negatively correlated with the amounts of acylsugar (Mandal et al., 2020).

### Expression of two *ISOPROPYLMALATE SYNTHASE* (*IPMS*) genes and acylsugar acyl chain profiles are altered in VIGS plants

IPMS catalyzes the first committed step in Leu biosynthesis, and there are five *IPMS* genes annotated in the *S. pennellii* genome (*Sopen06g018460, Sopen08g005060, Sopen08g005140, Sopen08g005150*, and *Sopen10g006110*) (Bolger et al., 2014a). Among these, *Sopen08g005140* and *Sopen10g006110* were noticeably downregulated in VIGS plants (80% reduction in transcript levels; Figure 3A). They share 99% sequence identity at the protein level (closest paralogs according to phylogenetic analysis of Ning et al. (2015)), and their promoters share 100% identity (although due to a gap in the genome sequence, we could extract only about 500-bp promoter sequence of *Sopen10g006110*). This indicates that a recent duplication event gave rise to these two sequences, including *cis*-regulatory elements. Among *IPMS* genes, these two members also exhibit noticeably high levels of trichome-enriched expression (Supplemental Figure 3). Together, these results indicate that ASTF1 directly or indirectly regulate expression of *Sopen08g005140* and *Sopen10g006110* in trichomes.

In addition to *Sopen08g005140* and *Sopen10g006110*, we observed that *Sopen08g005060* shows differential gene expression between high- and low-acylsugar-producing accessions of *S. pennellii* (Mandal et al., 2020). However, *Sopen08g005060* is expressed at a much higher level than *Sopen08g005140* and *Sopen10g006110* in leaf samples (Mandal et al., 2020), and it does not exhibit trichome-enriched expression (Supplemental Figure 3). Therefore, it could be involved solely in primary metabolism of Leu biosynthesis, or it could be a “junction gene” between primary and secondary (specialized) metabolism. On the other hand, *Sopen08g005140*, ortholog of *S. lycopersicum* trichome-tip-cell-expressed *IPMS3*, is a critical determinant of acylsugar acyl chain phenotype (3-methylbutanoate to 2-methylpropanoate ratio) (Ning et al., 2015). We observed a significant change in this ratio in VIGS plants (Figure 3B). These results indicate that ASTF1 regulates expression of trichome-specific *IPMS* genes, but not primary metabolic *IPMS* genes, and thereby controls acylsugar acyl chain profile.

### Acylsugar (and trichome flavonoid) metabolic genes are strongly co-expressed with *ASTF1*

Using expression profiles of *S. pennellii* genes in 48 RNA-Seq samples (38 from Mandal et al. (2020) and 10 from control and VIGS plants in this study), we performed weighted gene correlation network analysis (Langfelder and Horvath, 2008) to identify genes that are strongly co-expressed with *ASTF1*. We obtained 28 co-expressed modules, and *ASTF1* was placed in the “darkgrey” module with 125 other genes (Supplemental Data Set 3; Supplemental Figure 4, A and B). Module eigengene (first principal component) of the “darkgrey” module and expression profile of *ASTF1* showed strong correlation (Spearman’s rank correlation coefficient (SRCC)= 0.93; *P*= 6E-21), which indicated strong co-expression between *ASTF1* and other genes in this module. For example, *Sopen08g005140* (IPMS), *Sopen08g002520* (BETA-KETOACYL-ACP SYNTHASE III), *Sopen12g004230*-*Sopen12g004240* (BETA-KETOACYL-ACP SYNTHASE IV/II-like), *Sopen07g023200* (ACYLSUGAR ACYL-COA SYNTHETASE 1), *Sopen07g023250* (ACYLSUGAR ENOYL-COA HYDRATASE 1), *Sopen12g002290* (ASAT1), *Sopen04g006140* (ASAT2), and *Sopen11g026960* (ASAT3), which are all involved in acylsugar biosynthesis, as well as previously mentioned flavonoid metabolic genes, showed strong co-expression with *ASTF1* (SRCC= 0.89 to 0.96; *P*= 3.55E-17 to 2.46E-26). “Gene significance” for *ASTF1* and module membership for the “darkgrey” module genes also showed robust correlation (SRCC= 0.86; *P*= 4.9E-38) (Supplemental Figure 4C), which confirmed strong co-expression between *ASTF1* and acylsugar-flavonoid metabolic genes in the “darkgrey” module (Figure 3C).

### Identification of common motifs in promoters of potential ASTF1-target genes

Of the 125 genes that are strongly co-expressed with *ASTF1* in the “darkgrey” module, 34 were downregulated in VIGS plants. This suggests that expression of these 34 genes is regulated by ASTF1. Using the *de novo* motif discovery tool MEME (Bailey and Elkan, 1994), we identified three enriched motifs in the promoters of these 34 genes-GGWGGAKR, CACYCTCC, and AAATCGCTGY (e-value= 2.7E-11, 2.2E-04, and 1.5E-03, respectively), which are present in 20, 22, and 2 promoters, respectively (Figure 3D; Supplemental Data Set 4). The CACYCTCC and AAATCGCTGY motifs do not show significant similarities with known plant TF binding sites in the JASPAR CORE non-redundant database (Castro-Mondragon et al., 2021). However, the GGWGGAKR motif, which is present in promoters of several known acylsugar biosynthetic genes, returned matches with binding sites of nine TFs, all of which contain AP2 domains. For example, the top two hits are DREB2A and ERF055 (Supplemental Figure 5), which are involved in dehydration (Liu et al., 1998) and defense responses (Iwase et al., 2021), respectively, in *Arabidopsis thaliana*. These results suggest that genes strongly co-expressed with *ASTF1* and are downregulated in VIGS plants are regulated by AP2-domain containing TF(s), such as ASTF1. Multiple sequence alignment of the AP2 domain from ASTF1 and its homologs in the Solanaceae confirmed the presence of highly conserved DNA-contacting residues, validating ASTF1 as a bona fide AP2-family TF (Figure 3E).

### The orthologs of ASTF1 in *Nicotiana benthamiana* are also involved in acylsugar biosynthesis

The high level of sequence divergence for ASTF1 (Figure 1C) may also indicate a *S. pennellii*-specific function, rather than a Solanaceae-specific function. Therefore, we decided to determine if ASTF1 orthologs have a conserved function of acylsugar biosynthesis in other members of the Solanaceae. First, we performed a phylogenetic analysis, and using data of Moghe et al. (2017) (gene expression data in isolated trichomes and underlying tissues from four species-*S. nigrum, S. quitoense, Hyoscyamus niger*, and *Salpiglossis sinuata*), we identified two clades (called “upper” and “lower” hereafter) of ASTF1 homologs that showed trichome-enriched expression (Figure 4; Supplemental Figure 6). Next, we selected *N. benthamiana* for functional validation based on two facts-its phylogenetic distance from *S. pennellii*, and it being a well-known model for VIGS. BLAST identified three close homologs of ASTF1 in *N. benthamiana* (*Niben101Scf08050g00012, Niben101Scf02720g08001*, and *Niben101Scf03045g14001*), with all three of them showing similar RNA-Seq coverage in expression data (Bombarely et al., 2012). *Niben101Scf08050g00012* and *Niben101Scf02720g08001*, which form a monophyletic “upper” clade with *ASTF1*, share 95% sequence similarity at the nucleotide level with each other. We used one VIGS construct targeting both *Niben101Scf08050g00012* and *Niben101Scf02720g08001*, and VIGS resulted in 15% reduction in acylsugar production (3%-30% reduction in individual acylsugars; Figure 4B).

**Figure 4.**
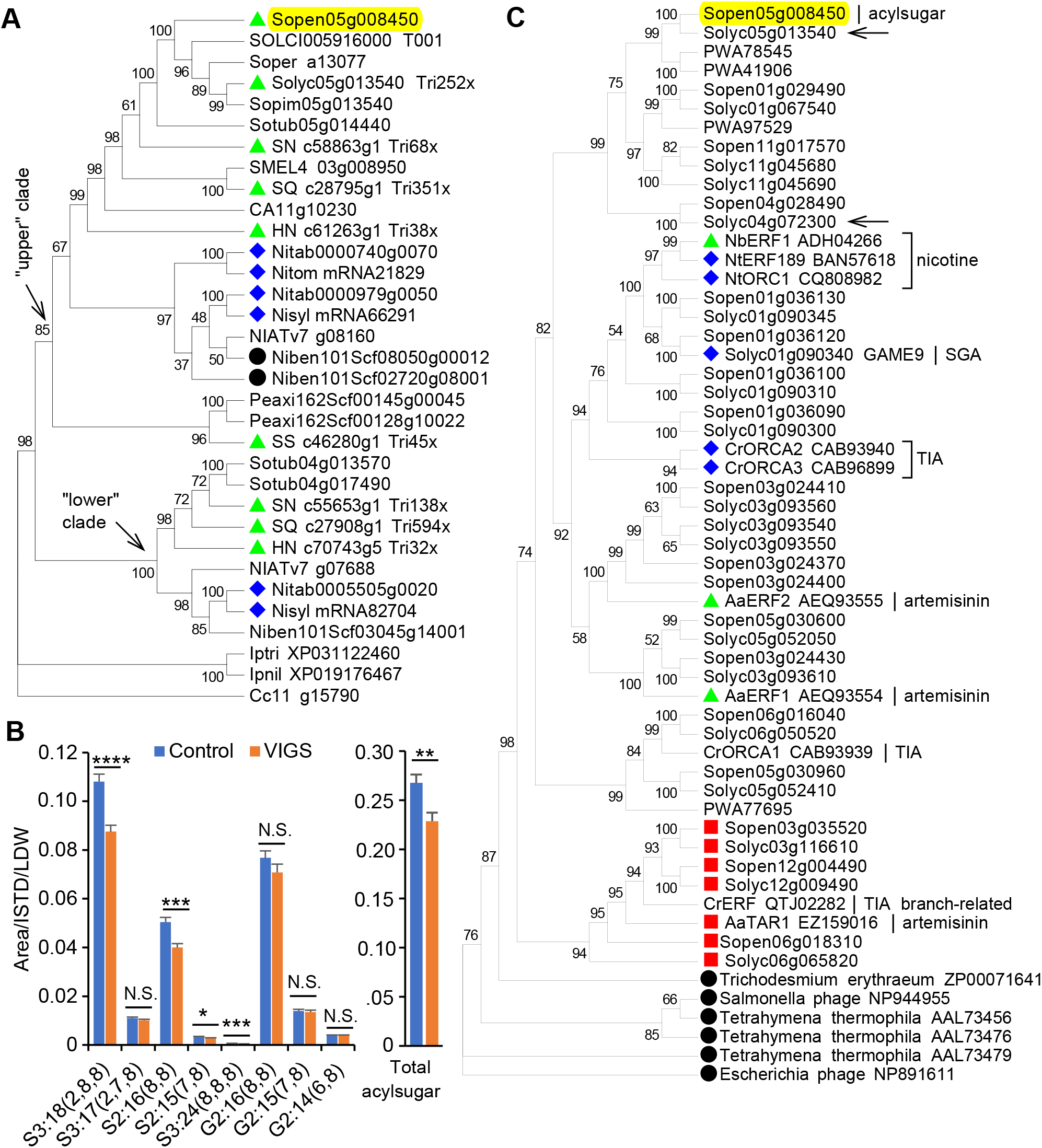
Phylogenetic analysis of ASTF1. (A) Maximum-likelihood based phylogenetic tree (topology) of ASTF1 (highlighted in yellow) and related sequences in the Solanaceae. Three non-solanaceous species are also included [Ipnil and Iptri= *Ipomoea nil* and *I. triloba*, respectively (Convolvulaceae; XP numbers indicate NCBI accessions); Cc= *Coffea canephora* (Rubiaceae)]. Bootstrap support values from 1000 replicates are shown. Members with available trichome-enriched expression data [(Ning et al. (2015) for *S. lycopersicum* and Moghe et al. (2017) for *S. nigrum, S. quitoense, Hyoscyamus niger*, and *Salpiglossis sinuata*] are marked with green triangles (Tri252x indicates 252-fold higher expression in isolated trichomes compared to underlying tissues). Homologs from *Nicotiana tabacum* and its two parents (*N. sylvestris* and *N. tomentosiformis*) are marked with blue diamonds. Two *N. benthamiana* sequences that were targeted for VIGS are marked with black circles. Sopen= *Solanum pennellii*; Solyc= *S. lycopersicum*; Sopim= *S. pimpinellifolium*; SOLCI= *S. chilense*; Soper= *S. peruvianum*; Sotub= *S. tuberosum*; SMEL= *S. melongena*; SN= *S. nigrum*; SQ= *S. quitoense*; CA= *Capsicum annuum*; HN= *Hyoscyamus niger*; NIAT= *Nicotiana attenuata*; Niben= *N. benthamiana*; Nitab= *N. tabacum*; Nisyl= *N. sylvestris*; Nitom= *N. tomentosiformis*; Peaxi= *Petunia axillaris*; SS= *Salpiglossis sinuata*. (B) VIGS in *Nicotiana benthamiana*. S and G indicate sucrose and glucose, respectively. Numbers followed by S or G indicate number of acyl chains and combined chain lengths, respectively; numbers within parentheses indicate individual acyl chain lengths; for example, S3:17 (2,7,8) indicates acylsucrose with three acyl chains and a combined chain length of C17 (C2, C7, and C8). Combined amounts of major acylsugars in control and VIGS plants are shown next to individual acylsugar bar graphs. Error bars indicate standard error (*n* = 11 biological replicates; N.S. *P* > 0.05, * *P* < 0.05, ** *P* < 0.01, *** *P* < 0.001, **** *P* < 0.0001; Welch *t*-test). (C) Maximum-likelihood based phylogenetic tree (topology) of ASTF1 and other specialized metabolic AP2/ERF members. Bootstrap support values from 1000 replicates are shown. Arrows indicate auxin-inducible members. Green triangles indicate jasmonate-inducible members. Blue diamonds indicate members that are both jasmonate-inducible and are present in tandem AP2/ERF TF clusters in the genome. Red squares indicate intron-containing members (information for CrERF is unknown); all other members in *S. pennellii* and *S. lycopersicum* do not have introns. Black circles indicate AP2 domains from non-plant proteins (cyanobacterium *Trichodesmium erythraeum*, ciliate *Tetrahymena thermophila*, and two viruses); these were included to infer possible evolutionary direction, based on information from Magnani et al. (2004). TIA= terpenoid indole alkaloid; SGA= steroidal glycoalkaloid. Aa= *Artemisia annua*; Cr= *Catharanthus roseus*; Nb= *N. benthamiana*; Nt= *N. tabacum*. GenBank accession numbers are given next to sequences, wherever applicable. PWA numbers indicate NCBI accessions for *Artemisia annua* AP2 members. Similar trees were obtained with neighbor-joining methods using only the AP2 domain from these sequences (Supplemental Figures 8 and 9).

It is worth noting that “lower” clade members showed more divergence in the AP2 domain sequence than “upper” clade members (Supplemental Figure 6). Additionally, there is an apparent loss of “lower” clade members in several species, especially in the tomato clade, and we observed signature of pseudogenization during potato-tomato divergence (Supplemental Figure 7).

### Phylogenetic relationships among specialized metabolic AP2/ERFs

AP2/ERF family TFs are key regulators of biosynthesis of several classes of specialized metabolites. Examples include GLYCOALKALOID METABOLISM 9 (GAME9) for steroidal glycoalkaloids (SGAs) in *S. lycopersicum* (Cardenas et al., 2016), NtERF189 and NbERF1 for nicotine in *N. tabacum* and *N. benthamiana*, respectively (Shoji et al., 2010; Todd et al., 2010), <u>o</u>ctadecanoid-derivative <u>r</u>esponsive *<u>C</u>atharanthus* <u>A</u>P2-domain (ORCA) proteins for terpenoid indole alkaloids (TIAs) in *Catharanthus roseus* (Menke et al., 1999; van der Fits and Memelink, 2000), and AaERF1-ERF2 for artemisinin in *Artemisia annua* (Yu et al., 2012; Tan et al., 2015). Phylogenetic analysis suggested that ASTF1 diverged from TFs that are critical regulators of these specialized metabolic pathways (Figure 4C; Supplemental Figures 8 and 9). One major distinction is that *ASTF1* is not part of a TF gene cluster (no AP2/ERF member found within 100 genes upstream and downstream of *ASTF1*), whereas many of these other specialized metabolic TFs are present in tandem AP2/ERF gene clusters on the genome. Most of these TFs are also jasmonate-inducible (for example, *ORCA2* and *ORCA3*, but not *ORCA1*). On the other hand, *Solyc05g013540* and *Solyc04g072300*, which are phylogenetically closer to *ASTF1* than jasmonate-inducible AP2/ERF members, have been reported to be auxin-inducible (Gupta et al., 2013; Capua and Eshed, 2017).

### Leaf initiation or trichome specialized metabolism or both

The ortholog of *ASTF1*/*Sopen05g008450* in *S. lycopersicum* (*LEAFLESS*/*Solyc05g013540*) has been reported to coordinate leaf initiation at the flanks of shoot apical meristem, with transient expression at incipient leaf primordia (expression stops after leaf initiation) (Capua and Eshed, 2017). Using data mining (Capua and Eshed (2017) for genes that are differentially expressed between wild-type plants and *leafless* mutants at the shoot apices, and Ning et al. (2015) for expression profiles of *S. lycopersicum* genes in stem trichomes and shaved stems), we observed that most of the genes that are downregulated in *leafless* shoot apices exhibit high levels of trichome-enriched expression, whereas most upregulated genes do not (Figure 5, A and B). In fact, more than 62% of the upregulated genes have much lower expression levels in stem trichomes compared to shaved stems. Additionally, we identified *S. pennellii*-*S. lycopersicum* (*Sopen*-*Solyc*) putative ortholog pairs that are differentially expressed in both this study and Capua and Eshed (2017). Of the 88 differentially expressed *S. pennellii* genes between control and *ASTF1*-VIGS plants, at least 22 genes’ putative orthologs in *S. lycopersicum* are also differentially expressed between wild-type and *leafless* shoot apices (Supplemental Data Set 5). Some gene pairs are missing because of missing and misannotations (for example, genes encoding MYRICETIN *O*-METHYLTRANSFERASEs; see Kim et al. (2014b)) and also due to limitations of reciprocal BLAST in identifying orthologous pairs. 13 gene pairs (including known acylsugar and flavonoid metabolic genes) were downregulated in both conditions (*ASTF1*-VIGS and *leafless*), and these genes exhibit noticeable trichome-enriched expression in *S. lycopersicum* (average 366-fold; Figure 5C). Together, these results indicate that both ASTF1- and LEAFLESS-regulated genes are preferentially expressed in trichomes and include acylsugar and flavonoid metabolic genes. Expression analysis of *ASTF1*/*Sopen05g008450* in different tissues of *S. pennellii* corroborated its trichome-preferential expression (Figure 5D), consistent with its role in trichome acylsugar biosynthesis.

**Figure 5.**
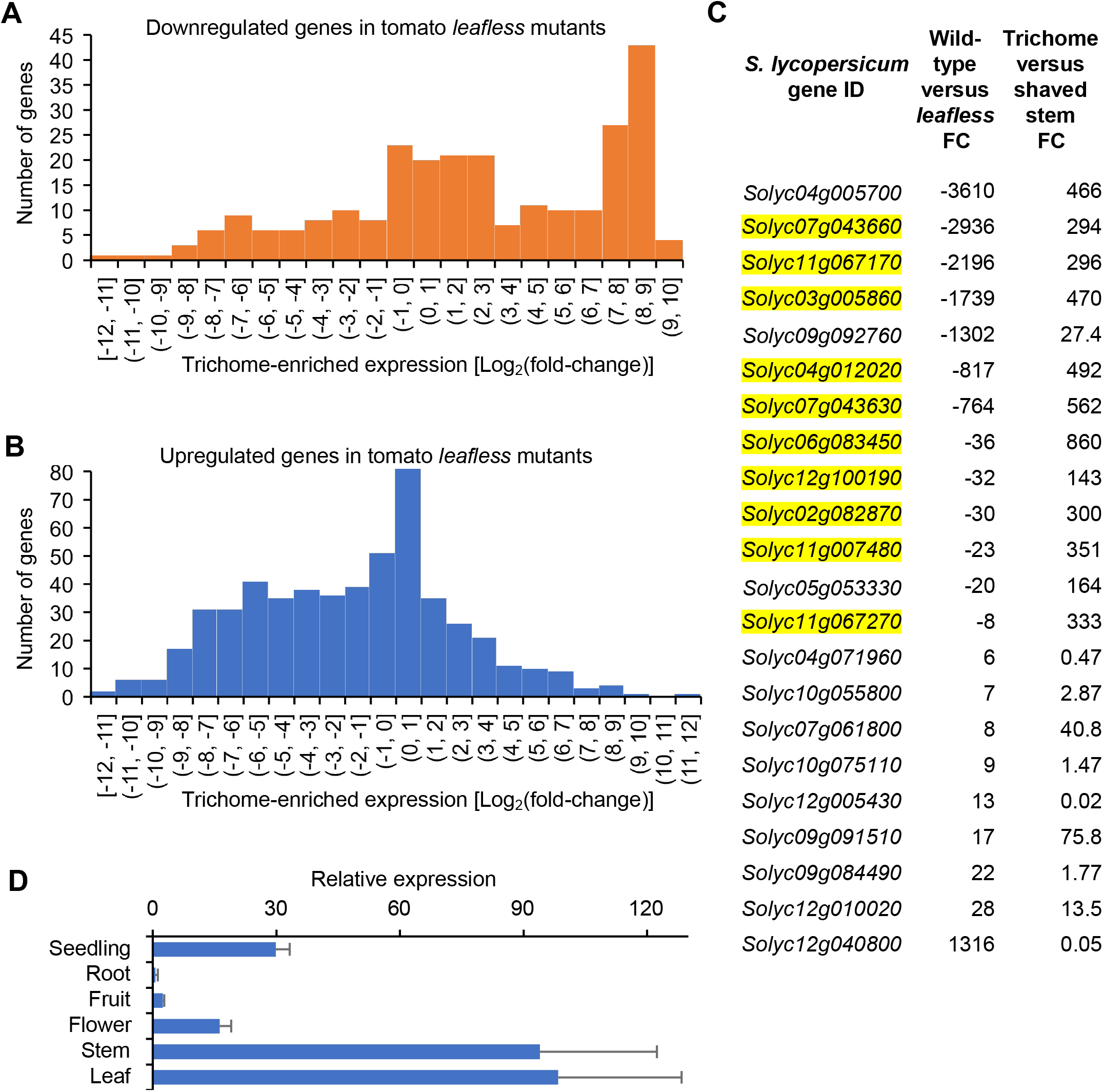
Comparing ASTF1 with LEAFLESS. (A and B) Histograms showing distribution of trichome-enriched expression (stem trichomes versus shaved stems) of genes that were downregulated (A) or upregulated (B) in *leafless* mutants’ shoot apices compared to wild-type *S. lycopersicum* shoot apices. Positive and negative log_2_(fold-change) values indicate higher and lower expression, respectively, in stem trichomes compared to shaved stems. (C) 22 *S. pennellii*-*S. lycopersicum* putative ortholog pairs that are differentially expressed in both this study and Capua and Eshed (2017). Only *S. lycopersicum* genes are shown (*S. pennellii* genes, annotations, and related information are given in Supplemental Data Set 5). Numbers indicate fold-change (FC) values. Negative sign indicates downregulation in *leafless* mutants. Known and candidate genes involved in acylsugar and flavonoid metabolism are highlighted in yellow. Data of Capua and Eshed (2017) and Ning et al. (2015) were mined to generate Figures 5A, 5B, and 5C. (D) Relative expression of *ASTF1* in different tissues of *S. pennellii*, as measured by RT-qPCR. Expression in leaf was set to 100%. Expression was detected in seedlings and predominantly in trichome-containing aerial tissues. Error bars indicate standard deviation (*n* = 3 biological replicates).

## DISCUSSION

In recent years, many enzymes involved in acylsugar metabolism have been identified, and these studies have also presented the opportunity to investigate the evolution of metabolic novelty and variation in acylsugar phenotypes conferred by structural genes (Ning et al., 2015; Schilmiller et al., 2015; Schilmiller et al., 2016; Moghe et al., 2017; Leong et al., 2019; Fan et al., 2020; Lou et al., 2021). However, regulatory factors involved in acylsugar metabolism remain elusive. Here, we report that ASTF1 is required for acylsugar biosynthesis and discuss regulation and molecular evolution of acylsugar production.

### Role of ASTF1 in acylsugar metabolism

In contrast to evolutionary conserved primary metabolites, specialized metabolites are restricted to specific taxonomic groups and specific tissue types. We used this knowledge to analyze trichome-enriched expression and sequence homology for each of the six candidate acylsugar metabolic TFs. Results from these analyses (Figure 1) indicated a role of ASTF1 in *S. pennellii* trichome acylsugar metabolism, and we used VIGS for functional validation of *ASTF1*. Both liquid chromatography- and gas chromatography-mass spectrometry analyses of leaf surface metabolites from control and VIGS plants confirmed this role (Figure 2). Next, we used RNA-Seq to show that many genes with known and putative roles in trichome specialized metabolism (acylsugar and flavonoid) were downregulated in VIGS plants (Figure 3A; Supplemental Data Set 1), indicating a role of ASTF1 in transcriptional regulation of these genes. This is consistent with downregulation of both acylsugar and flavonoid metabolic genes in response to imazapyr treatment (Mandal et al., 2020) and robust co-expression of these genes with *ASTF1* (Figure 3C). Acylsugars and flavonoids are structurally distinct compounds, and their building blocks are derived from different primary metabolic products; ASTF1 appears to integrate these two distinct specialized metabolic pathways in trichomes. Furthermore, promoter-sequence analysis of potential ASTF1-target genes suggested that many acylsugar metabolic genes are regulated by AP2/ERF-family TF(s), such as ASTF1 (Figure 3D; Supplemental Figure 5). Further work is required to confirm that ASTF1 indeed binds to promoters of these acylsugar metabolic genes *in vivo*.

### ASTF1 orthologs in the Solanaceae

Trichome-preferential expression of *ASTF1* and its orthologs in several acylsugar-producing solanaceous species (Figure 4A) suggests a conserved role in trichome acylsugar metabolism. VIGS of *ASTF1* orthologs in *N. benthamiana* supported this role. However, the effect of VIGS on acylsugar production in *N. benthamiana* was less pronounced than that in *S. pennellii*. Incomplete silencing of *Niben101Scf02720g08001* due to mismatches with siRNA may be responsible for this less-dramatic VIGS effect. Alternatively, it may indicate the critical role of other TF(s) in *N. benthamiana* acylsugar biosynthesis or a redundant activity from *Niben101Scf03045g14001* (“lower” clade member). Nonetheless, our results together suggest that *ASTF1* and its solanaceous orthologs are trichome-preferentially-expressed, if not trichome-specific, genes with a conserved role in acylsugar biosynthesis.

Tobacco (*Nicotiana tabacum*) is an allotetraploid with *N. sylvestris* as the maternal genome donor and *N. tomentosiformis* as the paternal genome donor (Murad et al., 2002). Because of undetectable (Moghe et al., 2017) or extremely low levels of acylsugars in *N. sylvestris* (Kroumova et al., 2016), it has been proposed that the acylsugar trait in *N. tabacum* originated from *N. tomentosiformis*. (Kroumova et al., 2016). Phylogenetic relationships among ASTF1 homologs in *Nicotiana* species (in particular, the presence of *N. tomentosiformis* ASTF1 ortholog in the “upper” clade, but not in the “lower” clade) (Figure 4A) are in good agreement with acylsugar phenotypes in these three tobacco species.

ASTF1 and LEAFLESS: Both *ASTF1* (Figures 1A and 5D) and *LEAFLESS* (Ning et al., 2015) are preferentially expressed in trichomes of developed leaves and stems, and they regulate expression of multiple acylsugar metabolic genes (Figure 5C). The role of LEAFLESS in acylsugar biosynthesis in *S. lycopersicum* remains to be confirmed (this cannot be tested in the *leafless* mutant because of its severe developmental defects; Capua and Eshed, 2017). However, based on our phylogenetic analysis and VIGS results in *N. benthamiana* (Figure 4, A and B, respectively), it is likely that ASTF1 orthologs, including LEAFLESS, are involved in acylsugar biosynthesis. Taken together, these results indicate remarkable spatiotemporal transition in expression and function of this TF-from leaf initiation at the shoot apical meristem (with transient expression at incipient leaf primordia) to specialized metabolism of acylsugars in trichomes (with constitutive expression in trichomes).

### Evolution of ASTF1 and emergence of acylsugar phenotype

Fan et al. (2020) reported that syntenic regions containing acylsugar biosynthesis-related gene clusters on both chromosomes 7 and 12 are located next to SGA gene clusters in *S. lycopersicum*. SGA biosynthesis is regulated by GAME9 (Cardenas et al., 2016), which appears to share a common ancestor with ASTF1. This raises the question of coevolution between GAME9 and ASTF1 and their target genes. Similarly, phylogenetic analysis of Moghe et al. (2017) suggests that ASATs have diverged from two transferases involved in TIA biosynthesis in *Catharanthus roseus*. TIA biosynthesis is regulated by ORCA proteins, and our analysis indicates that ASTF1 diverged from ORCAs (Figure 4C). Interestingly, *ASTF1* is strongly co-expressed with *Sopen07g029120* (Figure 3C; Supplemental Data Set 3), and BLAST identifies this gene as a homolog of *STRICTOSIDINE SYNTHASE* from *Catharanthus roseus* (McKnight et al., 1990; McKnight et al., 1991). STRICTOSIDINE SYNTHASE is a key enzyme in TIA biosynthesis, and its expression is regulated by ORCA proteins (Menke et al., 1999; van der Fits and Memelink, 2000). Therefore, we propose that divergence of ASTF1 from GAME9-ORCAs has been accompanied by divergence of acylsugar biosynthetic structural genes from alkaloid biosynthetic structural genes.

### Correlation with previously described quantitative trait loci (QTLs)

Using intraspecific populations from two accessions of *S. pennellii* (LA0716 and LA1912), Blauth et al. (1998) identified six QTLs associated with acylsugar level, and the QTL on chromosome 5 appears to be located near *ASTF1*. Additionally, Bonierbale et al. (1994) used segregating progenies of *S. tuberosum* x *S. berthaultii* crosses to study QTLs associated with acylsugar/trichome-mediated insect resistance. They mapped a single recessive gene (termed *bdr* by the authors) controlling acylsugar level to the short arm of potato chromosome 5 (between *TG441* and *TG379* markers). *Sotub05g014440* (“upper” clade member) appears to be located in this region. Interestingly, the *TG379* marker is located on chromosome 3 in *S. melongena* and on chromosome 11 in *Capsicum annuum* (https://solgenomics.net/). Based on our phylogenetic analysis, orthologs of *ASTF1* in these two species are also located on chromosome 3 and 11, respectively (*SMEL4_03g008950* and *CA11g10230*; Figure 4A). Taken together, our work demonstrates that ASTF1 is a positive regulator of acylsugar biosynthesis and opens the possibility of metabolic engineering of acylsugar production in solanaceous crops, such as tomato, potato, eggplant, and tobacco.

## METHODS

### Plant materials and growth conditions

*Solanum pennellii* (LA0716) seeds were obtained from the C.M. Rick Tomato Genetics Resource Center (University of California, Davis). Seeds were surface sterilized with 1.2% (w/v) NaOCl for 20 minutes and germinated on soil. Plants were grown at 24°C day/ 20°C night temperature under 16-h photoperiod with 150 µMol m^-2^ s^-1^ photosynthetically active radiation and 75% humidity. Seeds of *Nicotiana benthamiana* were germinated directly on soil, and plants were grown under the same conditions.

### Measurement of trichome-enriched expression

Trichome-enriched expression of selected genes were measured with reverse transcription-quantitative PCR (RT-qPCR), as described in Mandal et al. (2020). RT-qPCR primers used in this study are given in Supplemental Table 1.

### BLAST analysis

Sequences of six candidate TFs and selected proteins were obtained from the *S. pennellii* annotated gene models (Bolger et al., 2014a). BLASTp was performed at the National Center for Biotechnology Information (NCBI) website with the following options: expect threshold= 1E-05; word size= 6; matrix= BLOSUM62; max target sequences= 10. For each sequence, total BLAST score of the top hit was divided by protein length to get the normalized BLAST score.

### Virus induced gene silencing (VIGS)

The Solanaceae Genomics Network VIGS tool (http://vigs.solgenomics.net/) was used to design VIGS constructs. In order to minimize the probability of off-target silencing, we ensured that VIGS constructs do not have any 19-nt (or longer) matches with other annotated genes. For *S. pennellii*, a 580-bp fragment (coding sequence: 257-836) was cloned into the pTRV2-LIC vector (Dong et al., 2007), in the antisense orientation, to target *ASTF1* (*Sopen05g008450*). Overnight-grown *Agrobacterium tumefaciens* (GV3101) cells carrying pTRV1 and pTRV2 vectors were resuspended in infiltration medium (10 mM MES, pH 5.5; 10 mM MgCl_2_; 200 µM acetosyringone) and then mixed in a 1:1 ratio with a final OD_600_ of 1. Infiltration was performed at the first true leaf stage with a needleless syringe. Mixture of pTRV1 and empty pTRV2-LIC vectors were used as controls. After agro-infiltration, plants were grown for six weeks before collecting metabolites. A phytoene desaturase gene (*Sopen03g041530*) was used as a positive control to monitor the progression of VIGS. For *Nicotiana benthamiana*, a 400-bp fragment was used to target both *Niben101Scf08050g00012*.*1* and *Niben101Scf02720g08001*.*1* (*Niben101Scf08050g00012*.*1* coding sequence: 765-1164; antisense orientation). Plants at the four-leaf stage were infiltrated and then grown for additional four weeks before analysis. VIGS primers are listed in Supplemental Table 1.

### Liquid chromatography–mass spectrometry (LC-MS)

In order to collect secreted acylsugars, similar-sized young leaves were dipped into extraction solvent [acetonitrile:isopropanol:water = 3:3:2 (v/v/v) containing 0.1% formic acid and 100 μM propyl 4-hydroxybenzoate as the internal standard] and then gently agitated for 2 minutes. Exudates were analyzed using Q Exactive Focus coupled with Ultimate 3000 RS LC unit (Thermo Scientific). Separation of acylsugars were performed on Acclaim 120 (2.1 × 150 mm; 3 µm) C18 column (Thermo Scientific). Exactive Series 2.8 SP1/Xcalibur 4.0 software was used for data acquisition and processing. Two different methods were implemented for the analyses of *S. pennellii* and *N. benthamiana* exudates. For *S. pennellii*, extracted samples were separated by injecting 10 µL of sample into the C18 column that was housed at 30°C. Mobile phase consisted of 0.1% formic acid (eluent A) and acetonitrile with 0.1% formic acid (eluent B). Flow rate was set at 300 µL/min with the following gradient: 0–3 min, 40% B; 3-23 min, 40-100% B; 23-28 min, hold 100% B; 28.1-31 min, hold 40% B. The Q Exactive Focus HESI source was operated in full MS in negative electrospray ionization (ESI) mode. The spray voltage was set to 3.3 kV, and the sheath gas and auxiliary gas flow rates were set to 35 and 10 arbitrary units, respectively. The transfer capillary temperature and the auxiliary gas heater temperature were held at 320°C and 350°C, respectively. The S-Lens RF level was set at 50 v. The mass resolution was tuned to 70000 FWHM at m/z 200.

For *N. benthamiana*, extracted samples were separated by injecting 10 µL of sample into C18 column that was housed at 40°C. Flow rate was set at 500 µL/min with the following gradient: 0–5 min, 25% B; 5-20 min, 25-95% B; 20-25 min, hold 95% B; 25.1-30 min, hold 25% B. The spray voltage was set to 3.3 kV, and the sheath gas and auxiliary gas flow rates were set to 35 and 8 arbitrary units, respectively. The transfer capillary temperature and the auxiliary gas heater temperature were held at 320°C and 300°C, respectively. The S-Lens RF level was set at 70 v.

Targeted MS/MS of acylsugars from *S. pennellii* and *N. benthamiana* was performed using parallel reaction monitoring (PRM) mode. Acylsugar relative abundances in *S. pennellii* were determined by normalizing total peak areas of all acylsugars with peak area of the internal standard and leaf dry weight. For *N. benthamiana*, acylsugar relative abundances were determined by normalizing peak area of each acylsugar extracted ion chromatogram with peak area of extracted ion chromatogram of the internal standard and leaf dry weight. In cases of isomers, two or more peak areas were combined. LC-MS related information is given in Supplemental Data Set 6.

### Gas chromatography–mass spectrometry (GC-MS)

Transesterification of acylsugars and subsequent quantification of acyl chains were performed with gas chromatography-mass spectrometry (GC–MS), as described in Mandal et al. (2020). 100 μM methyl myristate (Sigma-Aldrich) was used as the internal standard.

### RNA-Seq library preparation and sequencing

Total RNA was extracted from leaves of control and VIGS plants of *S. pennellii* using the RNAqueous Total RNA Isolation kit (ThermoFischer Scientific), followed by treatment with the TURBO DNA-free kit (ThermoFischer Scientific) to remove genomic DNA. RNA-Seq libraries were prepared from poly(A)-selected RNA samples using the NEXTFLEX Poly(A) Beads 2.0 and Rapid Directional RNA-Seq kit 2.0 (PerkinElmer) and then sequenced on the Illumina NovaSeq 6000 SP v1.5 (2×150) platform at the Texas A&M Genomics and Bioinformatics service center. Sequence cluster identification, quality prefiltering, base calling and uncertainty assessment were done in real time using Illumina’s NCS 1.0.2 and RFV 1.0.2 software with default parameter settings. Sequencer .cbcl basecall files were demultiplexed and formatted into .fastq files using bcl2fastq 2 2.19.0 script configureBclToFastq.pl. 30-45 million paired-end reads were generated for each sample (average ∼35 million reads; average Phred33 quality scores >35; >99.97% base call accuracy).

### Differential gene expression analysis

RNA-Seq raw reads were processed with Trimmomatic v0.38 (Bolger et al., 2014b) using the following options: ILLUMINACLIP: TruSeq3-PE.fa:2:30:10:8, SLIDINGWINDOW= 1:10, SLIDINGWINDOW= 4:20, MINLEN= 120. This method ensured that all the ambiguous “N” bases were removed, and approximately 90% of the reads were retained. Processed reads were mapped to the *S. pennellii* genome (Bolger et al., 2014a), using the following options of STAR (Dobin et al., 2013): --genomeSAindexNbases 13, --sjdbOverhang 149, --outSAMmapqUnique 255, --alignIntronMin 50, --alignIntronMax 50000, --alignMatesGapMax 900. Read counts for annotated genes were calculated with StringTie2 (Kovaka et al., 2019) (reverse RF strand). Differential gene expression analysis was performed with edgeR (Robinson et al., 2010) using “Trimmed Mean of M-values” (TMM) normalization method and robust settings. Genes with low expression levels were excluded (<1 CPM in at least five samples after length-adjustment), and genes with >1.5-fold expression differences and false discovery rate < 0.05 (Benjamini and Hochberg, 1995) were identified as differentially expressed.

### Gene co-expression network analysis

RNA-Seq gene expression data from 48 leaf samples of *S. pennellii* (38 from Mandal et al. (2020) and 10 in this study) were used to identify co-expressed genes using weighted gene correlation network analysis (WGCNA) (Langfelder and Horvath, 2008) with the following options: softPower = 14; network type = signed; method = spearman; dissTOM= 1 – TOM; distM= dissTOM; deepSplit= 2; minModuleSize= 50. For each module, a module eigengene (ME) was selected as the representative of gene expression profile based on principal component analysis. For each gene, a module membership was calculated as the Spearman’s rank correlation coefficient (SRCC) between expression profile of that gene and module eigengene. Gene significance values for each gene in the “darkgrey” module were calculated as the correlation (SRCC) between expression profile of a gene and expression profile of *Sopen05g008450* (*ASTF1*).

### Identification of enriched motifs in promoters

We defined promoters as 2 kb sequences upstream of the putative transcription start sites, based on *S. pennellii* gene models (Bolger et al., 2014a). Promoters of four genes (*Sopen08g005140, Sopen08g002520, Sopen06g034810*, and *Sopen06g034830*) had long stretches of ambiguous “N” nucleotides (gaps in genome sequencing), which were replaced with three “N” nucleotides to reduce ambiguity in shuffled sequences. Enriched motifs were identified with MEME (Bailey and Elkan, 1994) using expect value < 1E-02 and the following options: -mod anr -minw 6 - maxw 10 -minsites 5 -maxsites 500. Enriched motifs were compared against JASPAR CORE (2022) non-redundant database of known plant TF binding sites (Castro-Mondragon et al., 2021). Motif (letter-probability matrix) comparison was performed with Tomtom tool (Gupta et al., 2007) using *Q* value (false discovery rate) threshold of 0.05 and the following options: -min-overlap 5 -dist pearson -thresh 0.05 -norc.

### Phylogenetic analysis

Sequences for Figure 4A were obtained as described below. Protein sequences of *Ipomoea nil* and *I. triloba* were obtained from NCBI. Protein sequences of *Coffea canephora* (v1.0) and the following solanaceous species were obtained from the Solanaceae Genomics Network (SGN): *Solanum pennellii* (v2.0), *S. lycopersicum* (ITAG release 4), *S. pimpinellifolium* (LA1589), *S. chilense, S. tuberosum* (ITAG release 4), *S. melongena* (v4.1), *Capsicum annuum* (CM334 v1.55), *Nicotiana attenuata* (v2), *N. benthamiana* (v1.0.1), *N. tabacum* (v4.5 edwards 2017), and *Petunia axillaris* (v1.6.2). BLASTn was used to identify *ASTF1* homologs in the following seven species: *S. peruvianum* (SGN *de novo* transcriptome), *N. sylvestris* (SGN mRNA), *N. tomentosiformis* (SGN mRNA), and the four species from Moghe et al. (2017) (*de novo* assembled transcriptomes of *S. nigrum, S. quitoense, Hyoscyamus niger*, and *Salpiglossis sinuata*). Nucleotide sequences of these seven species were translated in six possible frames (https://web.expasy.org/translate/) to obtain protein sequences with longest open reading frames. Careful observation revealed that *Hyoscyamus niger* sequence HNc61263_g1_i1 had a stretch of eight T nucleotides (coding sequence: 852-860), which caused a truncated protein sequence (probably an assembly artifact since c61263_g1_i2 has seven T nucleotides). Removal of one T restored the coding frame, which was in alignment agreement with other sequences, and it was used for further analysis. Protein sequences for Figure 4C were obtained from the Solanaceae Genomics Network (*S. pennellii* and *S. lycopersicum*) and the GenBank.

Multiple sequences for Figure 4A were aligned using MAFFT (Katoh and Standley, 2013) with BLOSUM62 matrix and the following options: mafft --localpair --maxiterate 1000 (gap extend penalty= 0.123; gap opening penalty= 1.53). Multiple sequences for Figure 4C were aligned with the following options: mafft --ep 0 --genafpair --maxiterate 1000 (gap extend penalty= 0.0; gap opening penalty= 1.53). Maximum-likelihood based phylogenetic analyses were performed with IQ-TREE 2 (Minh et al., 2020). Testing of the best substitution model [Jones-Taylor-Thornton (JTT) matrix (Jones et al., 1992) for Figure 4A and general “Variable Time” matrix (Muller and Vingron, 2000) for Figure 4C] was carried out using ModelFinder (Kalyaanamoorthy et al., 2017). Different models of protein evolution were compared, and the best models (JTT+F+I+G4 for Figure 4A and VT+F+I+G4 for Figure 4C) were selected based on lowest Bayesian Information Criterion (BIC) scores, Akaike Information Criterion (AIC) scores, and corrected Akaike Information Criterion (AICc) scores (all three methods gave uniform weights of 0.998 and 1 for Figure 4A and Figure 4C, respectively). Robinson-Foulds distance between maximum-likelihood tree and consensus tree (constructed from 1000 bootstrap trees) was zero for both Figure 4A and 4C. Neighbor-joining trees using only the AP2 domain from AP2/ERFs of Figure 4C were constructed with MEGA X (Kumar et al., 2018). Uniform rates among sites were used, and all positions containing gaps and missing data were completely deleted.

### Accession Numbers

Sequence data from this article can be found in the GenBank data libraries under the accession number OM811663 (ASTF1; Sopen05g008450). RNA-Seq reads used in this study were submitted to the NCBI Sequence Read Archive under the BioProject ID PRJNA821975.

## SUPPLEMENTAL DATA

**Supplemental Figure 1**. VIGS of ASTF1 in *Solanum pennellii*.

**Supplemental Figure 2**. Enriched gene ontology (GO) terms associated with differentially expressed genes between control and VIGS plants.

**Supplemental Figure 3**. Expression levels of *ISOPROPYLMALATE SYNTHASE* (*IPMS*) genes in isolated stem trichomes relative to underlying tissues of shaved stems.

**Supplemental Figure 4**. Weighted gene correlation network analysis (WGCNA).

**Supplemental Figure 5**. Comparisons between known plant TF binding sites in the JASPAR CORE non-redundant database and the GGWGGAKR motif.

**Supplemental Figure 6**. Multiple sequence alignment of ASTF1 and its homologs in the Solanaceae.

**Supplemental Figure 7**. Pseudogenization and gene loss of the “lower” clade members during potato-tomato divergence.

**Supplemental Figure 8**. Neighbor-joining phylogenetic trees using only the AP2 domain from specialized metabolic AP2/ERFs and their homologs.

**Supplemental Figure 9**. Multiple sequence alignment of the AP2 domain from specialized metabolic AP2/ERFs and their homologs.

**Supplemental Table 1**. List of primers used in this study.

**Supplemental Data Set 1**. Differentially expressed genes (DEGs) between control and VIGS plants.

**Supplemental Data Set 2**. Genes that were upregulated in VIGS plants.

**Supplemental Data Set 3**. Weighted gene correlation network analysis (WGCNA).

**Supplemental Data Set 4**. Enriched motifs identified in promoters of potential ASTF1-targets.

**Supplemental Data Set 5**. *S. pennellii*-*S. lycopersicum* putative ortholog pairs that are differentially expressed in both this study and Capua and Eshed (2017).

**Supplemental Data Set 6**. LC-MS results of acylsugars from *Solanum pennellii* and *Nicotiana benthamiana*.

## ACKNOWLEDGMENTS

We sincerely thank Wangming Ji for assistance with RT-qPCR and sample collections, Alan Pepper for helpful suggestions, Charlie Johnson and the staff of the Texas A&M Genomics and Bioinformatics center for performing Illumina sequencing, and the Texas A&M University High Performance Research Computer (especially Michael Dickens) for providing computational resources and assistance.

## AUTHOR CONTRIBUTIONS

SM designed and performed experiments, analyzed data, and wrote the manuscript. YHR performed chromatography–mass spectrometry analyses and reviewed the manuscript. TDM designed experiments, analyzed data, and wrote the manuscript. All authors read and approved the final manuscript.

